# Checkerboard synergistic data analysis using a Hill function-based approach

**DOI:** 10.1101/2025.02.05.636765

**Authors:** William G. Gutheil

## Abstract

This study presents a checkerboard data analysis approach using a Hill function (y = 1/(1+(x/K)^n^) to fit each column and row of checkerboard assay data. Column fits give a K (MIC) value in units of row concentration for each column antibiotic concentration (MIC_row vs [Col]), and row fits give an MIC_col value for each row antibiotic (MIC_row vs [Row]). Since the corresponding row and column concentrations are themselves MICs, this provides two sets of MIC vs MIC data pairs which can be plotted together as an isobologram. These MIC_A vs MIC_B values can be subjected to a second round of Hill function fitting, separately in x-y and y-x directions. Finally, a fit based on overlapping Hill functions is developed that allows x-y and y-x dimension fits to be performed simultaneously. Formula for fractional inhibitory concentrations (FICIs) as a function of fit parameters, and other features of these curves, are derived. This analysis also provides “n” (steepness) values from column and row fits, which are themselves dependent on the other antibiotic concentration and can be exceptionally, as in the case of ceftobiprole alone (n>10). This synergistic checkerboard analysis approach is implemented in MATLAB, which performs the fits and provides statistics variable values and alternative models significance.

## INTRODUCTION

Antimicrobial resistance (AMR) in pathogenic bacteria is a major public health threat (1-4). Given that the emergence of resistance to single agents has so far proven inevitable, methods to reverse or prevent the emergence of resistance, such as the development of antibacterial agent combinations, seem essential (5-7). Synergistic agent combinations, in which two agents show enhanced activity over either agent alone, are of obvious interest in such combinations (6, 8-11). Chemical library screening for synergistic agents has identified many such combinations in our own studies (12-15) and those of others (5, 6, 16-18). Some studies have also identified interesting three drug synergistic combinations (19-21).

The simplest approach to the analysis of checkerboard data is the “corners” method in which cells below a cutoff adjacent to cells above the cutoff are identified, and their row and column concentrations taken as the corresponding row and column MICs (Fig. 1A-B) (22). Plots of these values are known as isobolograms (23) (Fig. 1C). Isobolograms can be presented in their un-normalized form, as in Fig. 1C, or in their normalized form where the MICs are divided by their associated MIC0 (MIC of the corresponding pure compound) (Fig. 1B & D). Each data point in Fig. 1D will have an FIC_A (MIC_A/MIC0_A) and an FIC_B (MIC_B/MIC0_B), and a total FIC (∑FIC = FIC_A + FIC_B). The minimum of these ∑FICs is the FIC index (FICI) (24), which is generally considered indicative of synergy – a greater than simple additive effect of two agents – when it has a value less than 0.5 (25). More advanced analyses of synergetic interactions generally use fitting to global models of how the two agents interact (reviewed in (26-28)). Software based approaches for the analysis of drug synergies have been developed, including CompuSyn (29) and SynFinder (30-32).

**Figure 1:**
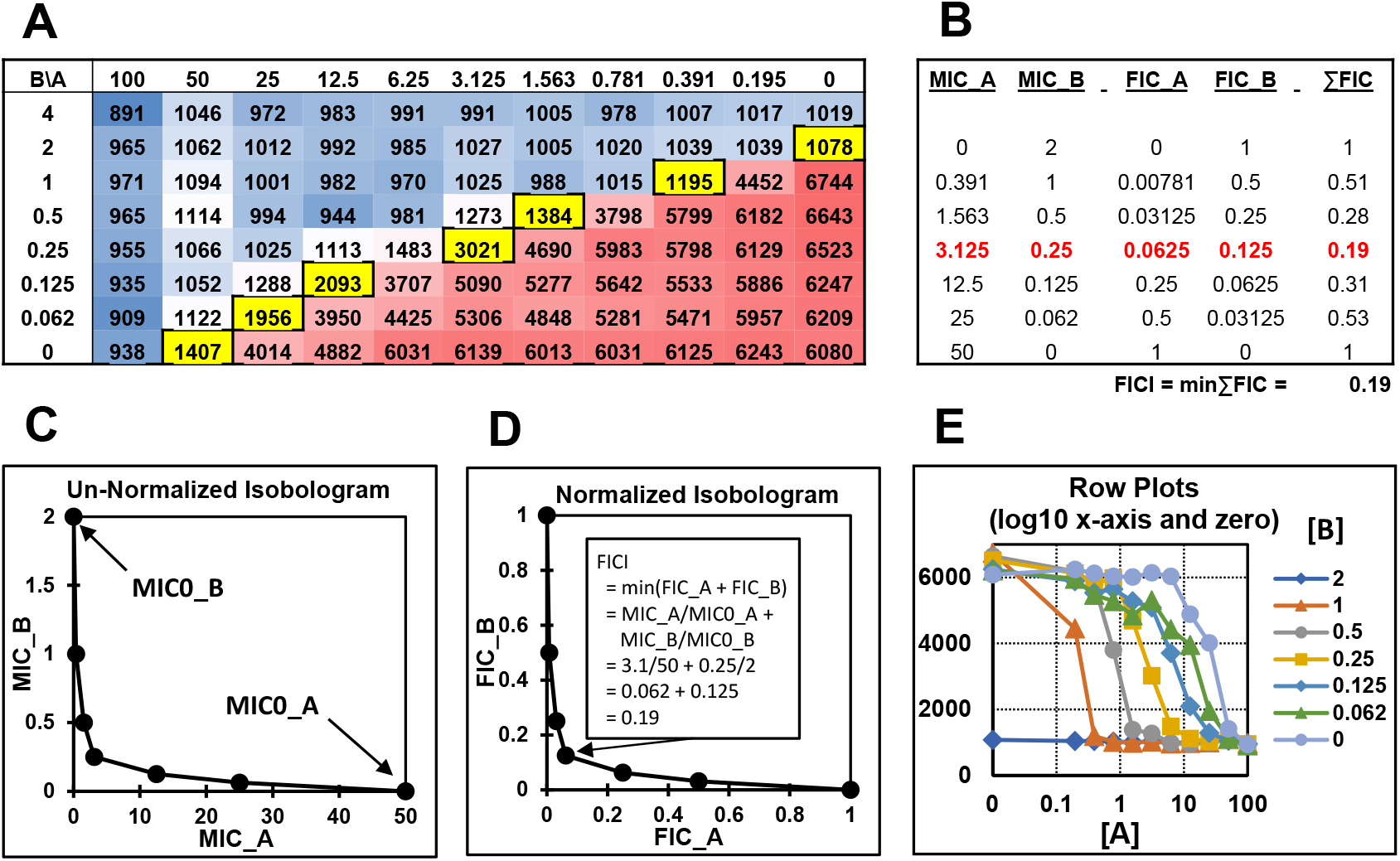
Checkerboard example data for a synergistic combination of cloxacillin (A=CLO in µM) vs ceftobiprole (B=BPR in µg/mL). Data from {Sharma, 2023 #161}. **A.** Color coded checkerboard data for [A] vs [B] effect on bacterial growth. Red = growth values at low [A] and [B], blue = growth values at high [A] and [B], yellow = “corner” values below midpoint cutoff (∼3500) used to provide MIC_A vs MIC_B values using the simple “corners” approach to checkerboard analysis. **B**. MIC values from panel A, and calculated FIC values, ∑FIC values, and the overall FICI value. Note that the rows in B are aligned with the yellowed entries in A. **C**. Corresponding un-normalized MIC_A vs MIC_B isobologram. FICI calculation for the lowest FICI data point shown in the inset. **D**. Normalized isobologram. **E**. Plots of row data from panel A showing cell growth signal vs variable [A] as a function of fixed [B]. A similar plot can be made for the columns in panel A.

Global fitting of checkerboard data has significant challenges in terms of complexity and interpretability. In the approach presented here an incremental approach is used. The first step is to treat each column and row in a checkerboard experiment as an individual MIC determination of one component against a fixed concentration of the other component. Fitting column data to a Hill function equation gives an MIC (midpoints) for the column in units of the row component vs the concentration of the column component (MIC_row v [col]), and row fits give MICs of the column component vs concentration of the row component (MIC_col v [row]). It can quickly be appreciated that these pairs of values are in fact MIC_row vs MIC_col and MIC_col vs MIC_row data pairs, and they can be plotted together to give data rich isobolograms. This also gives values of the exponential n term for each column and row fit, which are also functionally and mechanistically interesting since they vary across the columns and rows, i.e. they also vary with the fixed agent concentration.

In part 2 of this approach these MIC data pairs are fit again with a Hill-type equation in the x-y sense and then in the y-x sense. Mathematical relationships between variable, intercepts, and the FICI_x_, FICI_y_, and FICI are derived. This provides insight into the physical significance or the resulting fit parameter values. In part 3 a strategy to perform x-y and y-x fits simultaneously – bi-directional fitting – is developed and demonstrated. This incremental approach, particularly parts 1 and 2, are readily accessible. The bi-directional fitting approach in part three is more complex. A MATLAB program is provided with this approach implemented.

## METHODS and RESULTS

### Part 1: Fitting columns and rows for individual MICs

Example 8×11 checkerboard data for a synergistic combination is illustrated in Fig. 1A. This data was from a ceftobiprole (BPR) vs cloxacillin (CLO) checkerboard experiment originally obtained using 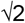 dilutions in 384 well plates (14). For easier visualization half of the columns and rows were deleted providing an 88 well data set with 2-fold dilution steps (Fig. 1A). This data was obtained using an Alamar Blue based assay procedure, but OD_600_ or any other cell growth reflective data can be analyzed the same way. The yellow entries and Fig. 1A are the 2D MICs identified using the simple corners approach (22). Fig. 1B summarizes the correspond MIC_A vs MIC_B pairs, the corresponding fractional FIC values, and their sums. The red entries identify the min∑FIC which is the FICI value characterizing the synergy (24). Fig. 1C shows the un-normailze isobologram plot of MICs, and Fig. 1D shows the normalized isobologram plot of FICs with the FICI point identified. Fig. 1E shows plots of the row values from Fig. 1A vs [A] at various B concentrations.

In other studies, we needed to analyzed large numbers of MICs from serial dilution data and found that fitting to a Hill-type equation worked well. Fitting with the hill function (aka E_max_, four parameter logistic regression) is widely used in many areas including for antibiotic effect data (33). The form used here is:

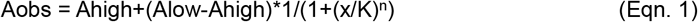

where Ahigh is the well value at high antibiotic concentration (no growth), Alow is the well value at low antibiotic concentration (full growth), x is the antibiotic concentration, K is the is the midpoint of the response curve in units of x, and n is a measure of “cooperativity” or hysteresis. Eqn. 1 could be used to fit each variable (Ahigh, Alow, K, and n) for each individual column or row in the checkerboard assay data (Fig. 1E type of data). However, Ahigh and Alow should be the same for the entire plate, and the fitting procedure used here therefore fits Ahigh and Alow globally over the entire plate, and K and n for each “fitable” column or row.

A MATLAB program (CBoard) has been developed to perform these fits (Supplemental zip file). Columns are fit first, followed by rows. This requires that fitable columns and rows be identified. To be fitable it is necessary for a column or row to have measured Aobs values both above and below the midpoint of the Aobs values. Otherwise, the MIC will be poorly or undefined by the data. For example, in Fig. 1E the flat curve for [B]=2 is not fitable. This fitability test also excludes the zero concentration values for columns and rows, since midpoints located between a low x value and a zero x value also give very poor fits for the K value. This is due to the K value not being bracketed by two concentration values, but by a value and zero. Column fit results are shown in Fig 2. A full summary of all fit results is provided in the CBoard output (Supplemental Information).

**Figure 2:**
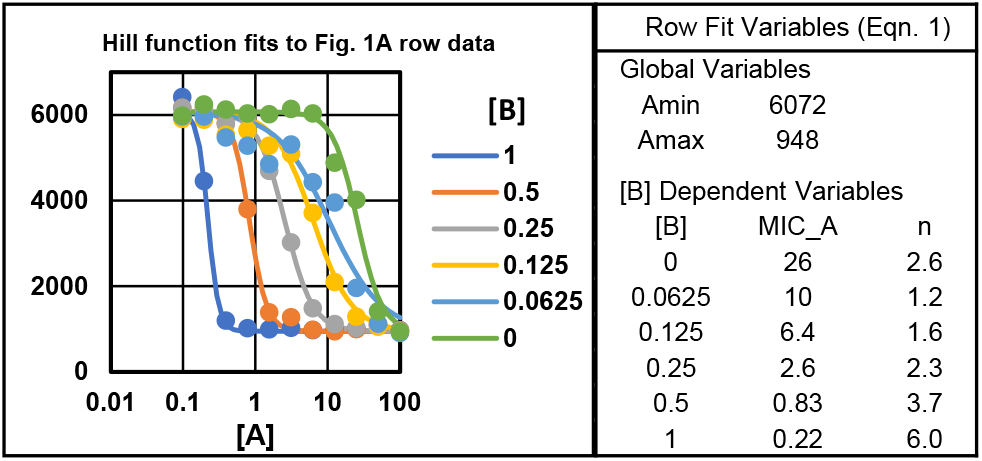
Column hill function fits for the MIC (K) and n values for cloxacillin (A = CLO in µM) vs ceftobiprole concentration (B = BPR in µg/mL).

Column fits and row fits give K and n values for each fitable column and row. The column fit K values are counted as the row MIC values, giving [col] vs MIC_row data pairs (will use [x] vs MIC_y_ herein for general purposes and A and B for the Fig 1 and 2 examples (Fig. 2A). Row fitting gives [y] vs MIC_x_ data pairs. Since the K values lie at the centered inflection point on the two-dimensional checkerboard data surface, the corresponding [x] and [y] values in these data pairs are also MICs in the perpendicular dimension. The result is a set of MIC_y_ vs MIC_x_ data pairs from the column fits and MIC_x_ vs MIC_y_ data pairs from the row fits. Note that these are not true MICs in the classical sense of the concentration necessary to reduce growth by 90%, however they are more tractable for mathematical data analysis. They can be converted into 90% inhibition MIC values mathematically (MIC_90% = K *x* 9^(1/n)^).

The CBoard program provides a series of graphics and reports to visualize and document these fits (Supplemental Figures S1-11, Reports 1-6). Fig. 2 shows an overlay of column fits. An overall column and row fit graphical summary is shown in Fig. 3. Plots of the fit MIC (K) values with error bars are shown in Figs. 3A-B. An extracted simple isobologram (corners method from Fig. 1A) is shown in Fig. 3C, and an overlay of panels 3A-C is shown in Fig. 3D. This illustrates that column and row fit MICs track with the simple isobologram values and provide many additional data points defining the synergistic antibiotic interaction. Fig. 3E and F panels show the dependence of n on variable concentrations of the two different agents. The n_BPR at [CLO] = 0 is high and is reflective of the very steep MIC curve for BPR at zero CLO (Fig. S3, right center panel).

**Figure 3:**
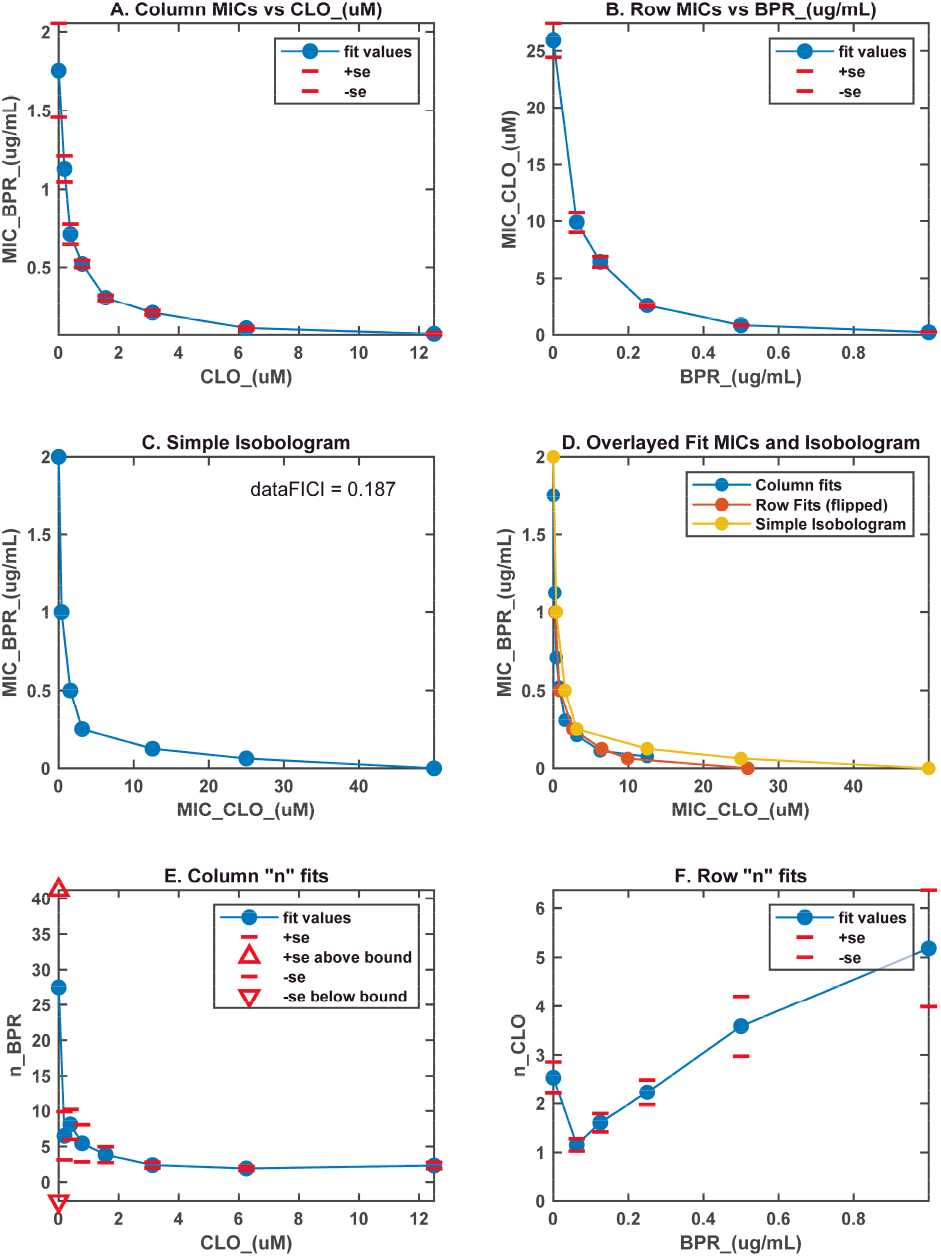
Graphical summary of column and row fits for BPR vs CLO checkerboard data. **A.** Column MIC fits. **B**. Row MIC fits. **C**. Simple isobologram. **D**. Overlay of column fits, row fits (x-y axes flipped), and simple isobologram. **E**. n values for BPR vs [CLO]. **D**. n values for CLO vs [CLO].

### Part 1: Mathematical Aspects

It is interesting to note that n in the hill function is directly related to the slope of these column and row curves at x = k, the midpoint of these curves. Eqn. 1 can be rewritten as:

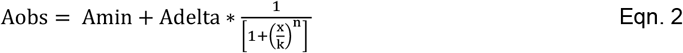

Differentiating with respect to x gives:

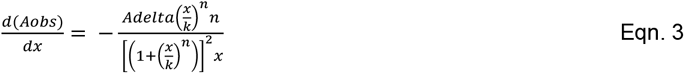

At x = k this reduces to:

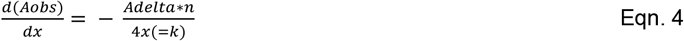

Moving k and A delta to the left side gives:

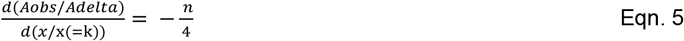

Demonstrating that the relative slope at the x = k midpoint is -n/4. When such curves are plotted on a logarithmic x axis – as in Figs. 1E and 2 – the apparent slopes of the curves will reflect this relationship. Plots of n vs [CLO] and [BPR] are shown Figure 3. For n > ∼ 5 the errors in n get quite large since the transition from uninhibited state to the inhibited state is very steep (Figs 2A for [B] = 1, and Figs. S2 and S3), and this uncertainty also translates to a lesser degree to uncertainty in the extracted MIC (k) values (Fig. 3A at [CLO] = 0). For this reason, we are now usually performing checkerboards using 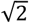 dilution steps.

### Part 2: Curve fitting of column and row MIC values

The MIC vs concentration (MIC vs MIC) curves in Fig. 3A-B also look like equilibrium binding isotherms, and therefore also amenable to fitting with a hill function, this time of the form:

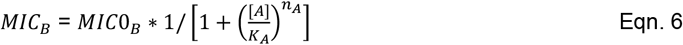

The data from Figs. 2A&B were therefore fit to Eqn. 6 (Fig. 4) with [A] = MIC_A. Initially it was unclear if the value of n in these fits would be substantially different than 1, so fits with n fixed to 1 (model 1; MDL_1) and n allowed to vary (model 2; MDL_2) were both performed. An F-test between the n=1 and n variable fits was then used to select the best model (34). The K values are readily interpretable as the midpoints of the observed effect of one agent on the other, and n-values again represent a slope factor. The results of these fits are also plotted and summarized in text files. Outlier identification flagged one point in Fig. 4C as an outlier. This same point caused the corresponding MDL_2 fit to be identified as best, rather than the simpler MDL_1. This is useful for illustrative purposes. Normally this single outlier would be removed and the data reprocessed for optimal results.

**Figure 4:**
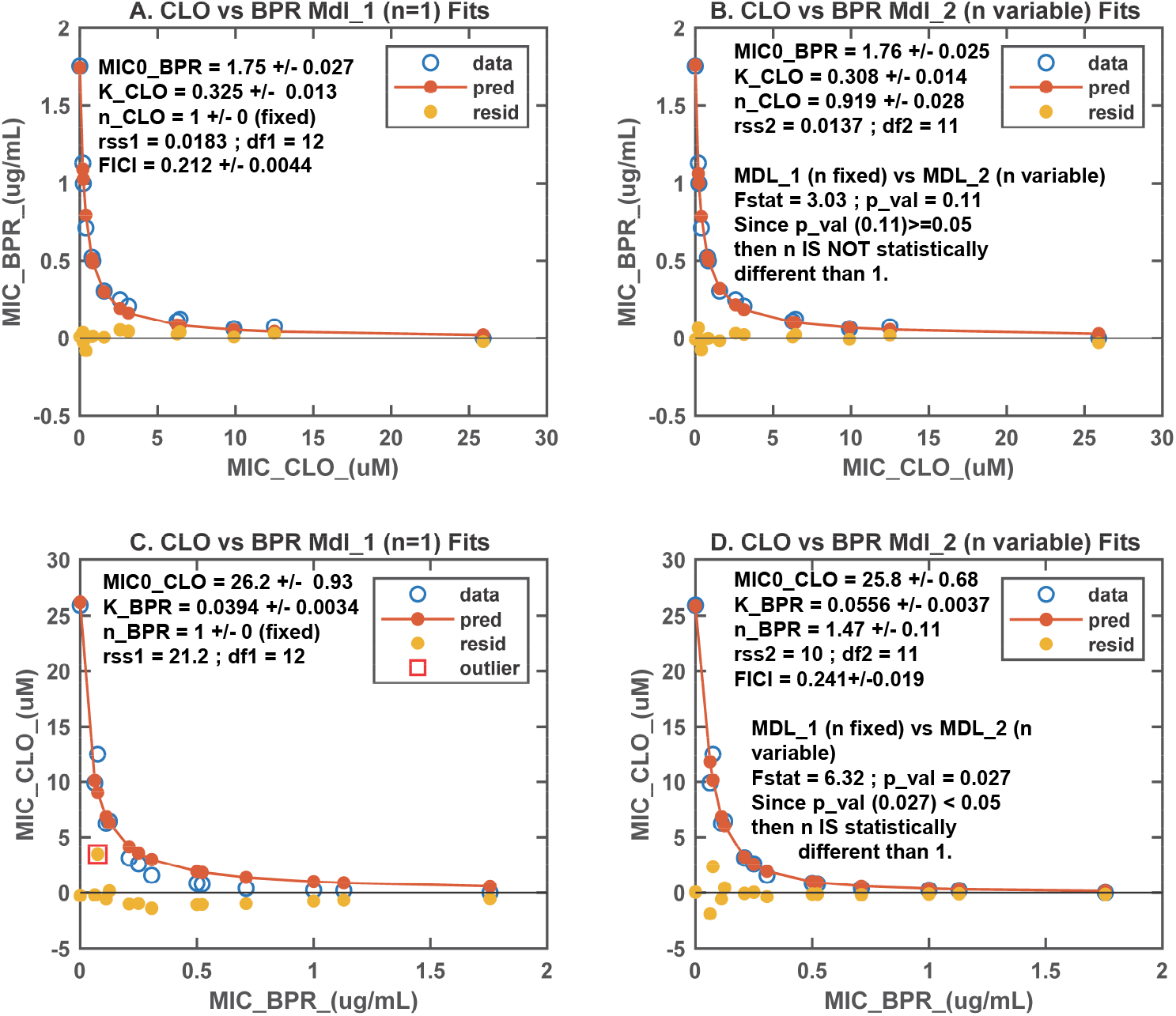
Graphical summary of MIC vs MIC fits for BPR vs CLO demonstration checkerboard data. **A.** CLO vs BPR MICs and fit with the n value fixed to 1 with fit variable values and SEs. **B**. The same with n variable. The improved fit with n variable is not significant. **C**. BPR vs CLO MICs and fit with the n value fixed to 1 (x-y axes flipped). In this figure an outlier is identified. **D**. The same with n variable. This time the reduction in rss is significant. However, this appears due to function contortion due to the outlier in panel C. The fit in panel C is also affected by the outlier. Statistical summaries included in the insets.

### Part 2: Mathematical aspects

This use of hill functions to model MIC_A vs MIC_B data in the form of un-normalized isobolograms (Fig. 4) allows for some mathematical insights to checkerboard assay behavior to be gleaned. Normalized isobolograms are more useful for this analysis and are the basis of the FICI measure of synergy (22, 24) (Fig. 1). This simply involves dividing all MIC_A values by MIC0_A to give FIC_A values, and all MIC_B values by MIC0_B to give FIC_B values (24). Normalized isobolograms are simply plots of FIC_A vs FIC_B.

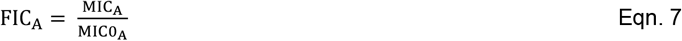

The FICs (∑FICs) are the sum of the FIC_A and FIC_B term for each data point.

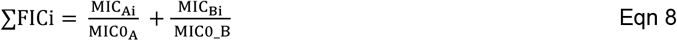

The minimum of these sums for synergistic combinations is taken as the FIC index (FICI, min∑FIC) (Fig. 1B). For synergistic combinations, the ∑FIC values vary from 1 at the x and y axes intercepts respectively down to a minimum value for synergistic agent combinations (Fig. 1C).

Eqn. 6 can be rewritten in normalized form in the following equivalent ways:

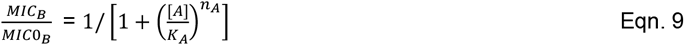

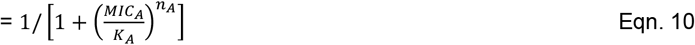

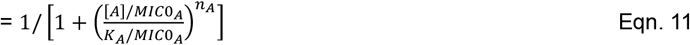

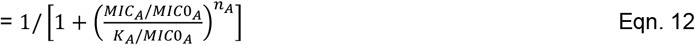

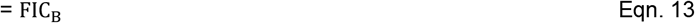

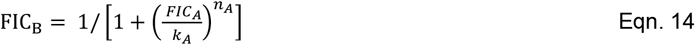

In the last of these the lowercase *k*_*A*_ is used to denote the normalized (by MIC0_A) *K*_*A*_ value. These formulae all have the general form of:

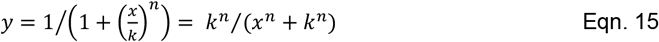

Eqn. 15 can be inverted to give:

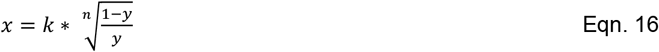

This last equation is the inverse hill function and allows axis inversion of Eqn. 15.

The FICI is the minimum of the following along the isobologram:

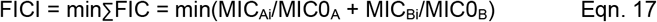

Considering the y term in Eqn. 15 as MIC_Bi_/MIC0_B_, the x term as MIC_Ai_/MIC0_A_, and the k term as K_A_/MIC0_A_, the ∑FIC formula can be written in terms of fit Hill function variables as:

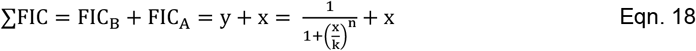

While this equation can be differentiated, finding the minimum (the FICI) by setting the differential equal to zero and solving for x is impractical algebraically, but can be accomplished by numerical minimization (this is implemented in the CBoard MATLAB program). However, if n = 1 Eqn. 15 reduces to the formula for a simple (non-cooperative) (normalized) equilibrium binding isotherm:

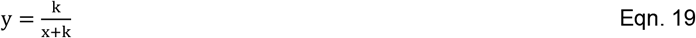

Eqn. 18 then reduces to:

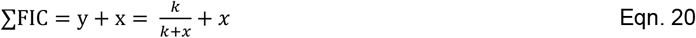

Differentiating Eqn. 20 with respect to x gives:

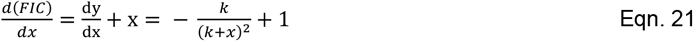

Setting the right side equal to zero and solving for x gives

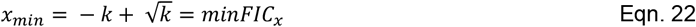

Which is the FIC_x_ value at the minimum FICI. Substituting this back into Eqn. 19 gives:

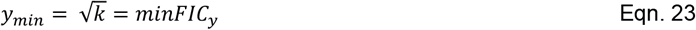

Given the FIC_x_ (Eqn. 22) and FIC_y_ (Eqn. 23) value at the minimum FICI. The FICI is then:

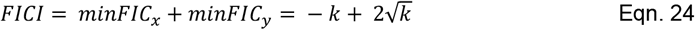

Note that as 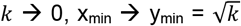,and 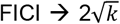.Starting with Eqn. 21, it can be shown that the (x_min_, y_min_) point is on the tangent line intersecting the y=k/(k+x) curve with a slope of -1. This in fact true of any function for FIC_y_ in terms of FIC_x_ since FICI = FIC_y_ + FIC_x_ = f(FIC_x_) + FIC_x_ and d(FICI)/d(FIC_x_) = f’(FIC_x_) + 1. Setting = 0 to find the minimum gives f’(FIC_x_) = -1.

There are a few other interesting features of normalized simple equilibrium (n=1) curves (Fig. 5). The point at which x=y on these curves is at:

**Figure 5:**
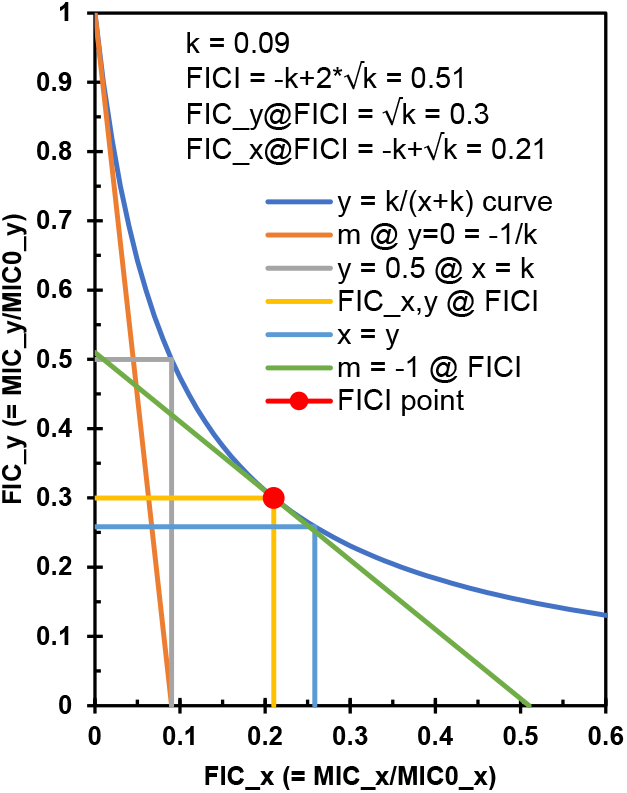
Illustration of relationships between k with a value of 0.09, FICs and FICI, and other characteristics for a normalized Hill function with n=1.

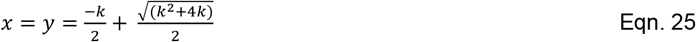

Another important feature of normalized hill functions is that the initial slope (Eqn. 14 and 19) at x = 0 is - 1/k, and this tangent line also intercepts the x axis at k. The k term (normalized apparent binding constant) reflects the relative sensitivity of FIC_y_ on x and vice versa. Finally, at x=1 in Fig. 4 there is a gap with the x-axis in hill function defined curves, when they should go to 0 at x=1, and this error can be shown to be described by:

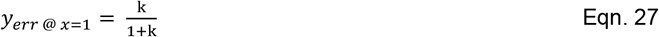

As k →0 this expression approaches k.

### Part 3: Bi-Directional Model Fitting

In Part 2 x-y and y-x fits are performed separately. It seems desirable to perform these two dimensions of fitting simultaneously. Most fitting efforts are based on the concept of an independent parameter (x value) determining the value of the dependent parameter (y value). Checkerboard MIC values are dependent variables in both the x-y and y-x dimensions. When plotted as MIC_x_ vs MIC_y_ values the different axes can have much different scales. Using normalized FIC_x_ vs FIC_y_ values avoids this disparity, and bi-directional model fitting was therefore performed on FIC transformed data using Part 2 MIC0 estimates.

As noted above, the hill function has a natural intercept at the (0,1) point on the FIC isobologram, but not on the (1,0) point. Overlaying the hill function from the x-y fit with the x-y axes swapped hill function from the y-x fit gives a good approximation over the full MIC/FIC plot range (Fig. 3D). This overlay can be defined in the x-y dimension using a combination of the hill function with x-y fit defined parameters and the inverse hill function (Eqn. 16) with y-x defined parameters:

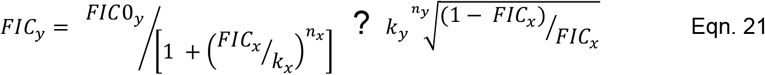

The ? denotes that some way of merging these two expressions is necessary. An identical equation can be written for predicting FIC_x_ values as a function of FIC_y_ values by swapping the “x” and “y” variables in this equation. Simply taking the average of these two terms in Eqn. 21 will not intercept the y- and x-axes at 1 as necessary. The first term intersects the y-axis at 1 when x is 0, and the second term intersects the x-axis at 1 when y is 0, so the first term needs to predominate at low x and the second at high x. Multiplying the first term by 1-x and the second by x and adding them together will work, but this weighting scheme does not match the underlying synergistic curves very well.

To derive a suitable weighting strategy a good but mathematically simple approximation to a synergistic isobologram curve is needed, one that passes through the points (0,1) and (1,0) and that has adjustable curvature. One such curve is:

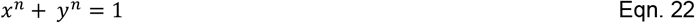

Which can be rewritten as:

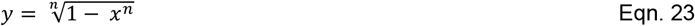

Such a curve can pass through any selected (x,y) point on a synergistic isobologram by appropriate choice of n. One such point might be the FICI point, but for strongly synergistic combinations this would dramatically emphasize very early points over later points. For the fitting algorithm used here the point selected was at average FIC_x_ and FIC_y_ component (FICI/2 effectively) of the FICI and 0.5 – the halfway point. This will always be above the FIC_y_ (or FIC_x_ point (except for weak synergies) and seems an appropriate intermediate mixing point keeping both terms in Eqn. 21 in play in the center of the isobologram. The value of n (N_FICI_) is determined numerically in this study. In the example data used the FICI ∼= 0.2, so ½ of this is 0.1, and for Eqn. 22 to pass through the (0.1, 0.5) point gives an N_FICI_ ∼ 0.53, which can be confirmed in Eqn. 22.

Eqn 21 can then be rewritten in the x-y direction as:

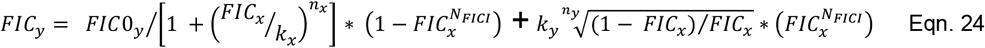

The converse (y-x) equation for FIC_x_ can be written in terms of FIC_y_ by swapping all the x & y subscripts.

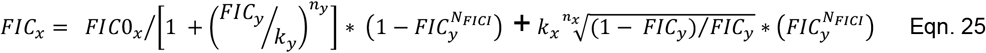

Eqn. 24 is compared against x-y data to generate an RSS_x_ value, and the converse relationship (Eqn. 25) against y-x data to generate an RSS_y_ value. These are added to provide an overall RSS measure of fit. The variable values are adjusted by to minimize this total RSS, providing a fit of both x-y and y-x variables against x-y and y-x data simultaneously. A few additional adjustments are necessary. When the ratio under the root in the second term is negative, this ratio is absolute valued to avoid a math error, and root assigned a negative value. This in essence projects values past FIC0_x_ into negative value space which enables good least squares fitting of experimental values slightly beyond the fit FIC0_x_ value. In this case the 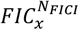 term is also set to 1. This root term is also undefined for FIC_x_ = 0 (in Eqn. 24), where this second term disappears due to the 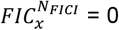 term, and this term is omitted when FIC_x_ = 0 in Eqn. 24 and when FIC_y_ = 0 in Eqn. 25. The accompanying MATLAB code in function H_fit_2D is annotated to describe these details.

Fitting to this model uses the Part 2 parameter value fits as a starting point. Statistical testing of n ≠ 1 is performed for both the n_x_ and n_y_ variable giving 4 models (Mdl_11 (n_x_, n_y_ = 1) Mdl_12 (n_x_ = 1), Mdl_21 (n_y_ = 1), and Mdl_22 (n_x_ and n_y_ both variable). An F-test is used for these incremental parameter statistical comparisons (34), and reports prepared. Supplementary information files contain a complete output from the analysis. Figure 6 shows the best bi-directional model to the example CLO vs BPR checkerboard data and some of the output fit variables. The MIC0 values, n values, and FICI values are comparable to the part 2 fits. (The K values in part 2 need to be divided by their corresponding MIC0 values to compare with the normalized k values fit in part3). Note that the x-y and y-x fits in part 3 are slightly different since even though the variable values are the same for both the mixing function weights them differently in each direction.

**Figure 6:**
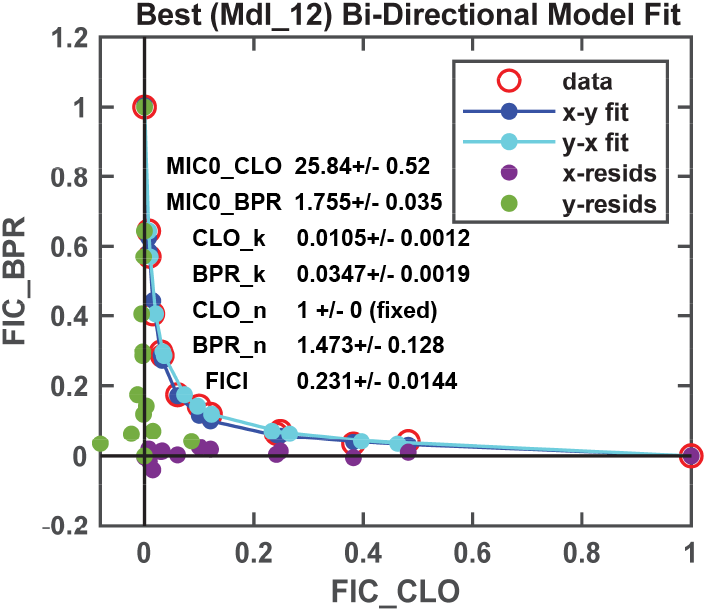
Bi-Directional fit to CLO vs BPR example data.

## Discussion

This effort was undertaken to allow data extraction from and parameterization of synergistic antibiotic combinations. The hill function was originally derived to describe cooperative equilibrium binding behavior in hemoglobin (35, 36) and has since found widespread use in many systems showing hysteresis in transitioning from one state (e.g. unbound) to another state (e.g. bound) (37). Other names for this function are the Emax and four parameter logistic function. This effort was divided into two main steps. The first major step is to fit column and row data with the hill function. This provides K and n values for each fitable column and row – e.g. column and row MICs vs column and row concentrations, which are in turn also MICs giving column and row MIC pairs. This approach is robust and provides a rich set of MIC vs MIC values for further analysis (Fig. 1D). The n values describe the steepness of the transition from the uninhibited to inhibited state (Fig. 3E & D). The n vs concentration dependencies are not considered further in this study. This appears a generally neglected feature of antibiotic action and synergistic interaction.

The second major step was to fit the resulting MIC vs MIC data with a second level of hill functions to assess how one agent affects the MIC of the other agent. Part 2 of this effort performed hill function fits to the resulting MIC_A vs MIC_B and MIC_B vs MIC_A data independently. Fits with n fixed to 1 (non-cooperative binding model) and n variable (cooperative binding model) were performed to determine if hysteresis (n∼=1) was significant. Outlier detection was also included in the part 2 analysis. In Part 3 of this analyses a bidirectional fitting procedure was developed using weighted overlays of the hill function and its inverse in both the x-y and y-x directions. This later approach works and provides a global hill function based fit to the data. Application of this approach to additional real checkerboard data will allow further development of this approach and further insight into the nature of synergistic interactions between antibiotics against target organisms. The use of the hill function for these fits provides physically interpretable fit variables for consideration and FICI values. This analysis approach is expected to facilitate mechanistic studies of synergistic antibiotic combinations.

## ASSOCIATED CONTENT

1. Supplemental Information file containing CBoard program output Figures 1-11 and output Reports 1-6.
2. A ZIP file containing a README file with instructions, 1 example raw data file for the main text example analysis, 8 additional example raw data files (14), example outputs, and the MATLAB CBoard processing scripts and functions.
3. The CBoard code and example files are also available via GitHub at: https://github.com/WGutheil/CBoard.git

## AUTHOR INFORMATION

Corresponding Author

*Phone: 816-235-2424

*

## ACKNOWLEDGEMENT

The author wishes to acknowledge Amar Deep Sharma for generating the original published data, and Vidit Minda for testing the CBoard program on new data. This work was supported by National Institutes of Health Grant R15 GM126502 (W.G.).

